# Slave Trade and Colonial Expansion Resulted in Strong Sex-Biased Admixture in South Africa

**DOI:** 10.1101/2023.09.06.556626

**Authors:** Haiko Schurz, Austin W. Reynolds, Gillian Meeks, Simon Gravel, Justin W. Myrick, Fernando L. Mendez, Cedric J. Werely, Paul D. van Helden, Eileen G. Hoal, G. David Poznik, Minju Kim, Caitlin Uren, Peter A. Underhill, Marlo Möller, Brenna M. Henn

**Author notes:** Co-first authors (Haiko Schurz, Austin W. Reynolds).

## Abstract

The colonial-period arrival of Europeans in southern Africa is associated with strong sex-biased migration by which male settlers displaced indigenous Khoekhoe and San men. Simultaneously, the importation of South Asian, Indonesian and Eastern African slaves may have contributed female-biased migration to Cape Town and surrounding areas. We examine the spatial and temporal spread of sex-biased migration from the Cape northward into Namaqualand and the southern Kalahari using genetic data from more than 1,400 individuals. In all regions, admixture patterns were sex-biased, with evidence of a greater male contribution of European ancestry and greater female contribution of Khoe-San ancestry. While admixture among Khoe-San, European, equatorial African, and Asian groups has likely been continuous from the founding of Cape Town to present-day, we find that Khoe-San groups further north experienced a single pulse of European admixture 6-8 generations ago. European admixture was followed by additional Khoe-San gene flow, potentially reflecting an aggregation of indigenous groups due to disruption by colonial interlopers. Male migration into the northern frontier territories was not a homogenous group of expanding Afrikaners and slaves. The Nama show evidence of distinct founder effects and derive 15% of their male lineages from Asian men, a pattern absent in the ≠Khomani San. Khoe-San ancestry from the paternal line is greatly diminished in populations from Cape Town, the Cederberg Mountains and Upington, but remains more frequent in self-identified ethnically indigenous groups. Strikingly, we estimate that Khoe-San Y-chromosomes were experiencing unprecedented population growth at the time of European arrival. Our findings shed light on the patterns of admixture and the population history of South Africa as the colonial frontier expanded.

## Author Summary

In this manuscript we report on whole Y-chromosome sequences and genome-wide data from over 1400 South Africans to answer questions surrounding the genetic impacts of European colonization in this region. We find that admixture patterns were sex-biased, with evidence of a greater male contribution of European ancestry and greater female contribution of Khoe-San ancestry across regions. But the patterns of admixture do vary regionally, for example, with the Nama deriving 15% of their Y-chromosome haplogroups from Asian lineages, a pattern absent in the ≠Khomani San. While estimates of admixture timing between Indigenous San and KhoeKhoe and European populations also varies geographically, they match closely with the first permanent European settlement in each region. Finally, we estimate that Khoe-San male lineages were experiencing unexpected population growth at the time of European arrival, while European male lineages experienced a founder’s effect consistent with previous research. Together these results provide a detailed look into the post-colonial genetic history of Southern Africa and illustrates the value of connecting genome-wide and uniparental markers to get a more complete understanding of population history in a region.

## Introduction

The Colonial period catalysed major social and genetic upheavals around the world. Many populations today can trace their genetic ancestry to historic admixture events between local autochthonous populations and European colonists. The population history and genetic architecture of admixed groups has been a major focus of genetic research in the Americas (Montinaro et al., 2015; Ongaro et al., 2019). Admixture patterns in the Americas are strongly sex-biased with European men contributing disproportionately in African Americans (Baharian et al., 2016; Bryc et al., 2015, 2010a; Mathias et al., 2016; Micheletti et al., 2020; Parra et al., 1998), Latinx, and Caribbean populations (Adhikari et al., 2017; Bryc et al., 2015, 2010b; Jordan et al., 2019; Moreno-Estrada et al., 2013; Ongaro et al., 2019). Correspondingly, females contributed greater African and Native American ancestry. These genetic patterns are corroborated by the extensive historic records of the colonial period in the Americas. Because of their complementarity with historic records, genetic techniques can be especially useful in contexts where written documents are relatively sparser with regards to human movements and interactions, such as in the case of rural regions of the Western and Northern Cape Provinces in South Africa.

Traditionally, sex-biased admixture has been estimated by comparing the proportion of uniparental markers such as mitochondrial DNA (mtDNA) and Y chromosomes that come from each ancestral population (Carvajal-Carmona et al., 2000; Mesa et al., 2000; Parra et al., 2001; Sans et al., 2002). A female bias is present when lineages from a specific ancestral population are increased in the mtDNA compared to the Y chromosome, while a male bias is present when there is an increased proportion of Y chromosome to mtDNA from a particular ancestral population (Rasteiro et al., 2012). An alternative method of investigating sex biased admixture is to analyse the ancestral distributions of the X chromosome compared to the autosome (Bryc et al., 2010b, 2010a; Wang et al., 2008). The X chromosome spends 2/3’s time in females where recombination occurs and 1/3 time in males without recombination. The autosomes, on the other hand, undergo approximately the same rate of recombination in both males and females. As a result, the X chromosome retains longer linkage disequilibrium blocks than the autosomes (Cox et al., 2010; Pereira et al., 2015). For example, in the case of a two-way admixed population, if the mean ancestral component of population A is higher on the X chromosome compared to the autosomes, then this is an indication of female sex bias from ancestral population A and male bias from ancestral population B. Alternatively, it could also indicate a male bias from both population A and B with more influx from population A (Goldberg and Rosenberg, 2015).

While prior work on sex-biased admixture in Africa has focused on the Bantu expansion across the Central rainforest and surrounding the Kalahari (Bajić et al., 2018; Destro-Bisol et al., 2004; Pierron et al., 2017), comparatively little has been written about the genetic consequences of European colonization and specifically, resulting sex-biased admixture patterns in Africa (Korunes et al., 2021). Perhaps the best studied population in this regard is the South African “Coloured” (SAC) population. Today, approximately 47.5% of the population in the Western Cape and 43.7% of those in the Northern Cape Provinces of South Africa self-identifies as “Coloured”. These communities are referred to as “Coloured” based on an artefact of the colonial and Apartheid-era South African terminology (ADHIKARI, 1996, 1992). Together, the SAC population numbers over 8 million individuals in South Africa today.

The SAC population has the most complex admixture history in South Africa, with ancestry derived from 5 primary sources (de Wit et al., 2010; Petersen et al., 2013; Quintana-Murci et al. 2010). The colonial period in southern African began when Dutch colonists settled in the Cape of Good Hope in 1652 and Europeans slowly expanded geographically over the next 250 years (Penn 2005). The Dutch East India Company and other traders brought slaves to the Cape from Madagascar, southeast Africa, South Asia and Indonesia numbering ∼63,000 individuals (Groenewald, 2010). Communities with multiple ancestries emerged in the Cape Colony, derived from the European colonists, imported slaves, as well as indigenous Khoekhoe laborers. The mode and timing of this admixture over the next few hundred years remains poorly described. As the colony expanded throughout the 18^th^ century, discrimination towards these mixed communities increased (Penn, 2005), resulting in individuals migrating towards the northern and eastern frontiers of the Cape colony. As they fled the expanding colony, they encountered other Khoe-San groups, and either became integrated with them (Penn, 2005) or formed independent communities (Edgar and Saunders, 1982; Nurse and Jenkins, 1975; Ross, 1975).

Khoe-San is a collective term for the indigenous peoples of southern Africa. Khoe-San communities harbour a high level of genetic diversity representing the earliest population divergence among human populations (Henn et al., 2016, 2011; Ragsdale et al. 2023). The diverse range of groups include both hunter-gatherers (“San”) and cattle/goat/sheep pastoralists (“Khoekhoe”) who share a common genetic ancestry (Uren et al., 2016). Their interaction with Bantu-speaking agro-pastoralist populations, who derive from western Africa, varies over time and space (Pickrell et al., 2012; Schlebusch et al., 2017; Vicente et al., 2019). Bantu-speaking groups largely displaced or incorporated Khoe-San populations in the eastern half of South Africa over the past 1,500 years but western Bantu-speakers only more recently expanded into Namibia and Botswana from Central/West Africa.

Much like the genetic admixture history of American populations, initial mtDNA and Y chromosome results for the SAC in Cape Town are broadly consistent with the detailed historical records of the Cape Colony (Elphick et al., 1989). However, the patterns of admixture and population history of SAC populations further from the Cape are still poorly understood. Furthermore, the historical records regarding the rural regions of the Western and Northern Cape Provinces are very limited, so genetic data provides an opportunity to learn more about not just when Europeans first began expanding into these regions, but when substantial interaction between colonists and Khoe-San groups begins as the colonial frontier expanded. In the present study, we use genome-wide SNP data and whole Y-chromosomes from five South African populations to address these questions. We reconstruct the timing and extent of admixture patterns and find a south/north gradient in admixture. We also test for patterns of sex-specific gene flow and infer recent changes in effective population size as a result of the historic patterns of movement and gene flow.

## Results

To examine the spatial and temporal spread of sex-biased admixture in South Africa, we generated genome-wide SNP data from 1434 individuals from 5 South African populations on the Illumina MEGA and H3Africa array platforms: the ≠Khomani San (*n* = 170), Nama (*n* = 110), SAC from the Cederberg (n=161), SAC from Upington (n=183), and SAC from Cape Town (*n* = 810). To better understand the dynamics of male-specific lineages in these regions, 272 African samples were short-read sequenced for 8.9Mb of the Y-chromosome capturing targets informative for haplogroup assignments (Poznik et al., 2013) (see Methods, Table S3).

### Global autosomal and X-chromosome ancestry inference

To resolve patterns of colonial sex-biased admixture and earlier migration in southern Africa we first contrasted autosomal and X chromosome ancestry distributions. Differences in the distributions of ancestral components between the autosome and X chromosome reflect the average of many independent loci. Following quality control and merging with appropriate ancestral populations, there were a total of 558,213 autosomal and 13,399 X chromosome variants in the dataset. Estimates for ancestry inference were performed using ADMIXTURE (v1.3); this version allows for X chromosome-specific admixture inference. The choice of reference populations was based on prior publications and with the aim of keeping the number of genotyped SNPs above half a million. For the SAC from Cape Town, source populations included European; equatorial African from Nigeria, Kenya, and South African Bantu-speakers; Namibian and South African Khoe-San; Indian; and Japanese and Chinese – as the Indonesian, Malagasy components have no comparable genomic datasets. Ancestry components in the Nama and ≠Khomani can be primarily attributed to three sources: Khoe-San, European, and Equatorial African, while the SAC also have a minority of their ancestry from recent South and East Asian sources (Fig. 1).

**Fig. 1:**
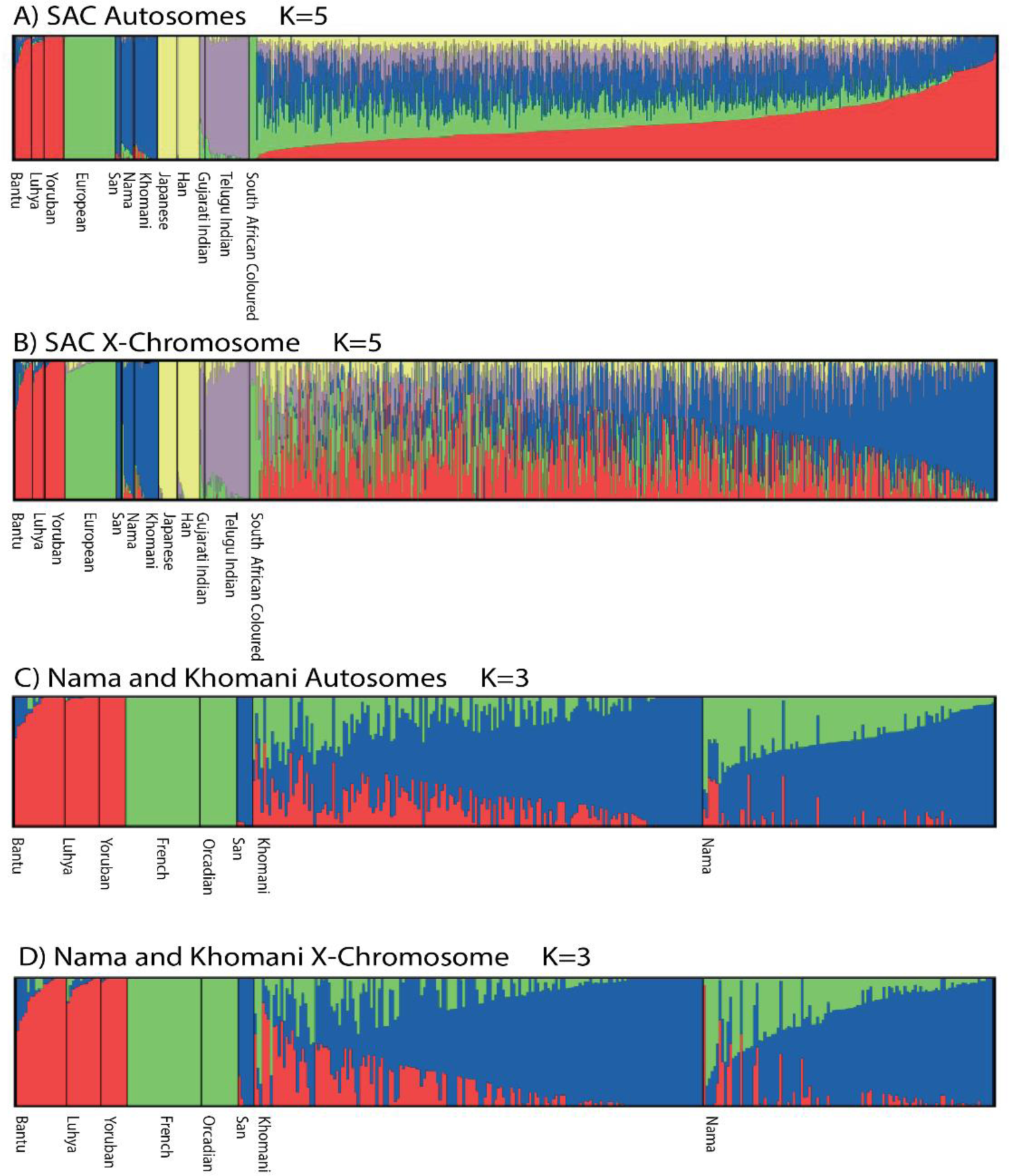
Admixture plot with reference populations for all SAC autosomes. (A) SAC X-chromosomes (B), Nama and ≠Khomani autosomes (C), and Nama and ≠Khomani San X-chromosomes (D). Each column represents the admixture distribution for an individual and the order of the individuals are the same for the autosome and X chromosome plots.

For the SAC from Cape Town, mean autosomal ancestral components are consistent with previous results of admixture analysis (Chimusa et al., 2013; de Wit et al., 2010; Daya et al., 2014). As shown in Table 1, the equatorial African (p-value = 7.16e^-5^) and European (p-value = 1.47e^-31^) components are significantly higher on the autosomes than the X chr, suggesting a male bias. Contributions from the indigenous Khoe-San population (p-value = 1.78e^-20^) are significantly higher on the X chromosome compared to the autosome, indicating a female bias. The East Asian component passed the significance threshold (p-value = 0.038) following multiple testing correction suggesting a female bias, while the South Asian component did not reach statistical significance (p-value = 0.32), but the mean values for the South Asian components were slightly higher in females compared to males.

**Table 1:** Sex-biased distribution of each ancestral component in the SAC, Nama and ≠Khomani San population.

| Population | Ancestry | Mean |  | Sex Bias | p-value | p-value adjusted |
| --- | --- | --- | --- | --- | --- | --- |
|  |  | Autosome | X chr |  |  |  |
| SAC<br>(Cape<br>Town) | Equatorial<br>African | 0.280 | 0.267 | Male | 1.43e<sup>-5</sup> | 7.16e<sup>-5</sup> |
|  | European | 0.205 | 0.162 | Male | 2.93e<sup>-32</sup> | 1.47e<sup>-31</sup> |
|  | Khoe-San | 0.271 | 0.328 | Female | 3.56e<sup>-21</sup> | 1.78e<sup>-20</sup> |
|  | South Asian | 0.147 | 0.148 | Female | 0.064 | 0.32 |
|  | East Asian | 0.096 | 0.099 | Female | 0.0076 | 0.038 |
| Nama | Damara | 0.0402 | 0.064 | Female | 0.0658 | 0.1975 |
|  | Khoe-San | 0.6663 | 0.712 | Female | 0.0226 | 0.0677 |
|  | European | 0.2935 | 0.224 | Male | 0.0002 | 0.0007 |
| #Khomani<br>San | Bantu-speaking<br>African | 0.155 | 0.152 | Male | 0.0829 | 0.2489 |
|  | Khoe-San | 0.704 | 0.755 | Female | 0.0063 | 0.0191 |
|  | European | 0.141 | 0.093 | Male | 0.0014 | 0.0042 |

The Nama show a significant signal of male biased admixture from the European population (p=0.0007) and the X/autosome ratio of the Khoe-San ancestry component is suggestive of female-biased admixture in the Nama, but does not maintain statistical significance after multiple-testing correction (p=0.0677) (Table 1 and Fig. 1). Unlike the SAC (Table 1), the Nama show no strong sex-biased influence in the equatorial African (likely Damara) ancestry component (p=0.1975). The ≠Khomani San show significant male biased admixture from the European population (p=0.0042) and significant female-biased admixture from the KhoeSan ancestry component (p=0.0191). Like the Nama, the ≠Khomani show no significant sex bias in the Bantu-speaking African ancestry component (p=0.249) (Table 1 and Fig. 1).

### Timing of Admixture in the Western and Northern Cape

To gain insight into the timing, order and magnitude of admixture events in the Nama and SAC populations, we modelled the distribution and decay of the local ancestry patterns using the TRACTs software (*Materials and Methods*). Comparable analysis for the ≠Khomani San data were published in Uren et al. (2016). Local ancestry inference (LAI) of phased array data was performed with RFMix.

The initial date of admixture among the primary source populations for the SAC is well defined, with a bound of 1652 at the time of documented European arrival. It is unclear whether gene flow between indigenous Khoe-San groups and Europeans occurred immediately, and when South Asian, Malagasy, equatorial African and Indonesian slaves came into contact with these groups. After partitioning SAC genomes into 4 putative ancestries, we first estimated the timing of admixture under a simple “two pulse” model. The ancestral population is seeded with all possible combinations of the 4 ancestries (Table S1), thus allowing for gene flow between two or more groups. Similarly, later pulses could involve any arbitrary combination of ancestries. We then successively added three and four pulses of gene flow. The highest likelihood fit represented a three pulse model with similar levels of gene flow from all groups (LL=-540; Fig. 2); similarly, the next best model (LL=-558) indicated 3 pulses of gene flow among all groups, followed by a 4th pulse of European, Equatorial Africa and Asian gene flow but without additional Khoe-San input. We caution, however, that these models did not always fit the data well, resulting in an excess of short tracts and paucity of long tracts. These model fits also sometimes resulted in admixture dates occurring slightly earlier than our expected bound of 1652 (i.e., ∼11-18 generations ago assuming 30-20 years per generation). Even under the best fit model, the initial pulse of admixture among all groups occurred 22 generations ago, which clearly predates the time of European arrival. We conclude that the data support generally continuous gene flow among all four ancestral groups dating back to the initial founding of Cape Town, but uncertainty in LAI or violations of model assumptions (i.e. random mating) likely prevents a more fine-grained estimate of varying admixture events over the past 300 years.

**Fig. 2:**
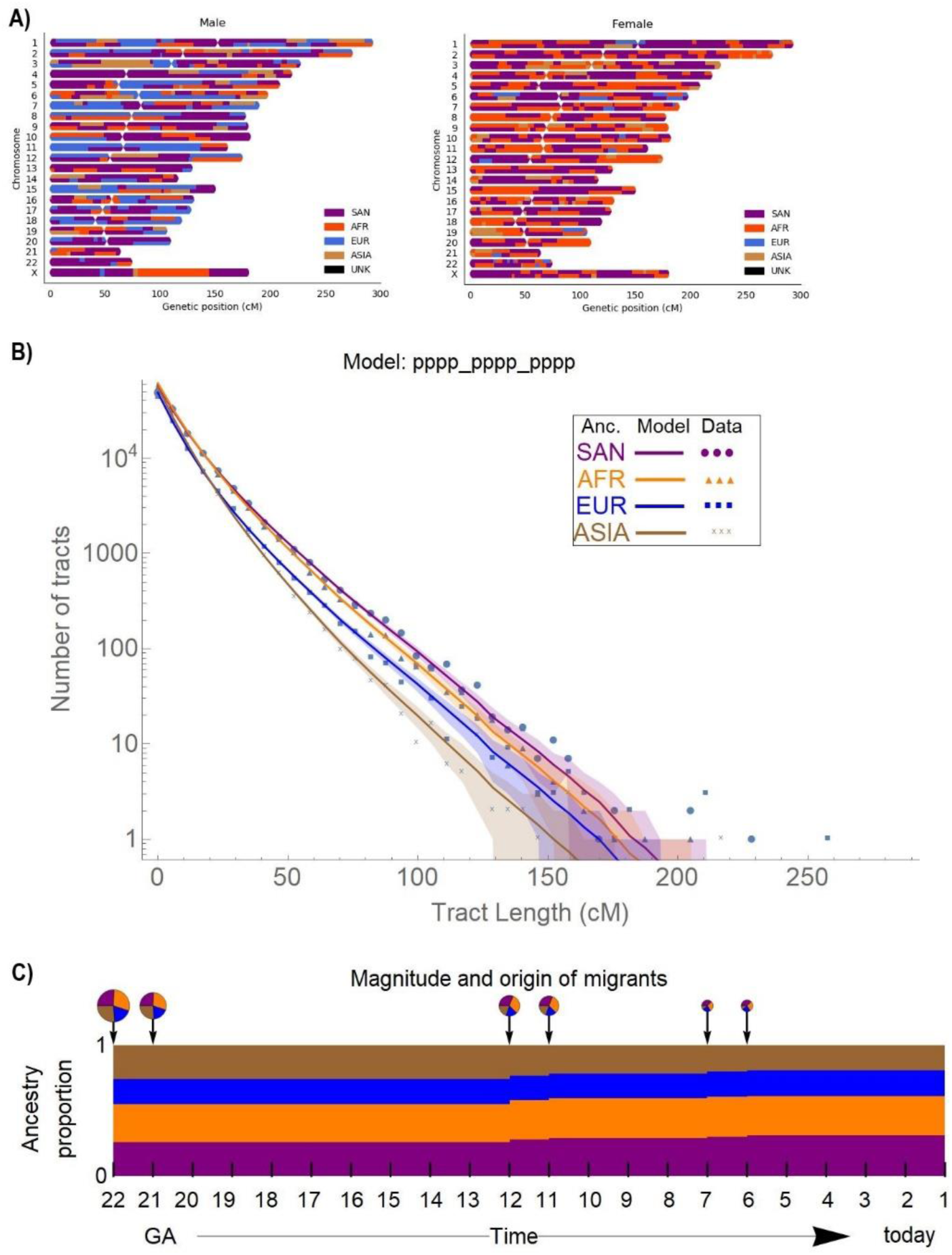
Local ancestry and TRACTs analysis for the SAC population showing. **A)** Local ancestry karyogram for a male (left) and female (right) representative of the 4-way admixed SAC population, made using RFMix (Maples et al., 2013). **B):** TRACTs decay results for the most likely 3-pulse model, pppp_pppp_pppp. Colored dots show the observed distribution of ancestry tracts for each ancestry, solid lines show the best fit from the most likely model, and shaded areas indicate confidence intervals corresponding to ±1 SD **C)** Estimation of migration time (GA: generations ago), volume of migrants (circle plots) and ancestry proportion over time (Gravel, 2012). San: KhoeSan, AFR: Equatorial African, EUR: European, ASIA: South and South East Asia.

Model fits for the Nama were better and more historically interpretable. The Nama community represented in this study presently live 800km north of the Cape. While colonial expeditions up the Atlantic coast occurred soon after arrival, with the aim of prospecting and trading for cattle, permanent settlements first occurred across the Orange River in 1790 (Penn 2005). We considered 3 possible ancestral sources for the Nama, with the same populations as for the SAC model but without the fourth (Asian) ancestral component. The TRACTs model with the highest likelihood suggests three gene flow events (model: ppx_xxp_ppx, Table S2, Fig. 3). Under this model, the Nama would have received gene flow from an equatorial African population between 26-27 generations (∼780-810 years) ago (Fig. 3). This is consistent with the timing of Khoe-San gene flow into northern Namibian Damara, Khwe and Tswana from Pickrell et al. (2012), but much older than estimates into the southern Kalahari groups like the ≠Khomani (Uren et al., 2016).

**Fig. 3:**
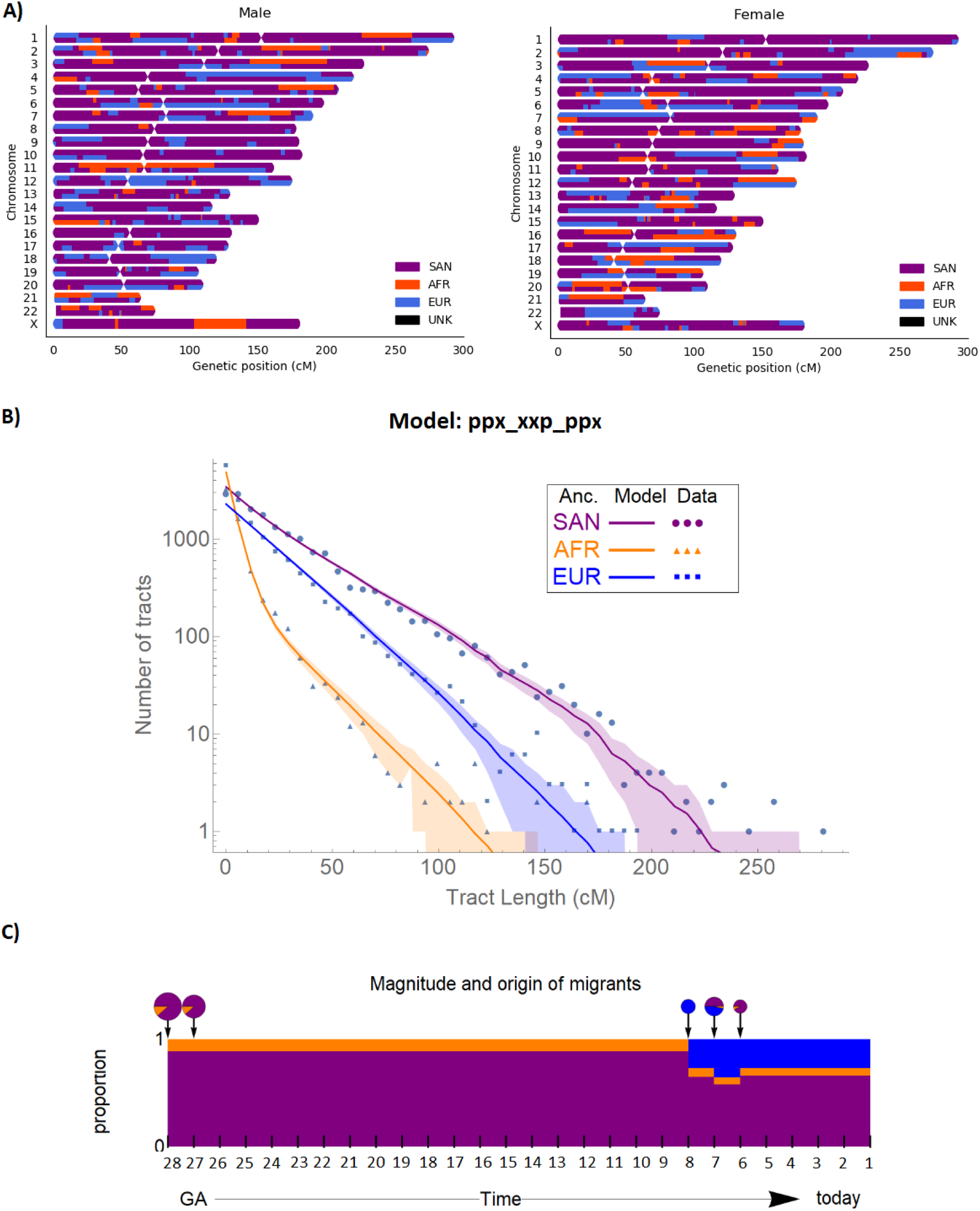
Local ancestry and TRACTs analysis for the Nama. **A)** Local ancestry karyogram for a male (left) and female (right) representative of admixed Nama, estimated from RFMix. **B)** TRACTs decay for the most likely 3-pulse model, ppx_xxp_ppx. Colored dots show the observed distribution of ancestry tracts for each ancestry, solid lines show the fit from the model, and shaded areas indicate confidence intervals corresponding to ±1 SD. **C)** Estimation of migration time (GA: generations ago), proportion of migrants (circle plots) and ancestry proportion over time. San: Khoe-San, AFR: Equatorial African (Damara), EUR: European.

European admixture into the Nama begins during the second pulse of the model, occurring between 7 and 8 generations (210 – 240 years) ago, consistent with the migration of Dutch farmers into Little Namaqualand (Fig. 3). Finally, we observe a recent pulse of Khoe-San ancestry between 6-7 generations (180-210 years) ago (Fig. 3). A similar pulse of European gene flow, quickly followed by Khoe-San gene flow is also observed in the ≠Khomani San population (Uren et al., 2016). This is also supported by our demographic reconstruction of Khoe-San male lineages (Fig. 5), which show a rebound in effective population size during the colonial period, consistent with the incorporation of multiple Khoe-San populations as they retreated from the expanding colonial frontier and annexation of tradtional Khoe-San land.

### Sources of Male-biased Gene Flow

We investigated male gene flow concordance between Y-chromosome results and autosomal and X-chromosome sex-biased admixture results (Table 1). For context, we determined Y-chromosome haplogroups for 584 individuals from southern African Khoe-San descendant populations, as well as 173 men from Zimbabwean Bantu-speaking farming groups, Central African forager groups, and Kenyan pastoralists. Haplogroups were inferred for the Cape Town SAC along with Coloured individuals from the Cederberg and Upington using SNAPPY version 0.2.1 (Severson et al., 2018) applied to Illumina SNP array data. Haplogroups for the 7 other populations were inferred from newly generated NGS Y-chromosome capture data using yhaplo (Poznick 2016). Here, we discuss Y-chromosome variation from Khoe-San, European, South Asian / Indonesian and eastern African ancestors; for a full description of haplogroups frequencies see Fig. 4 and Supplemental Table 3.

**Fig. 4:**
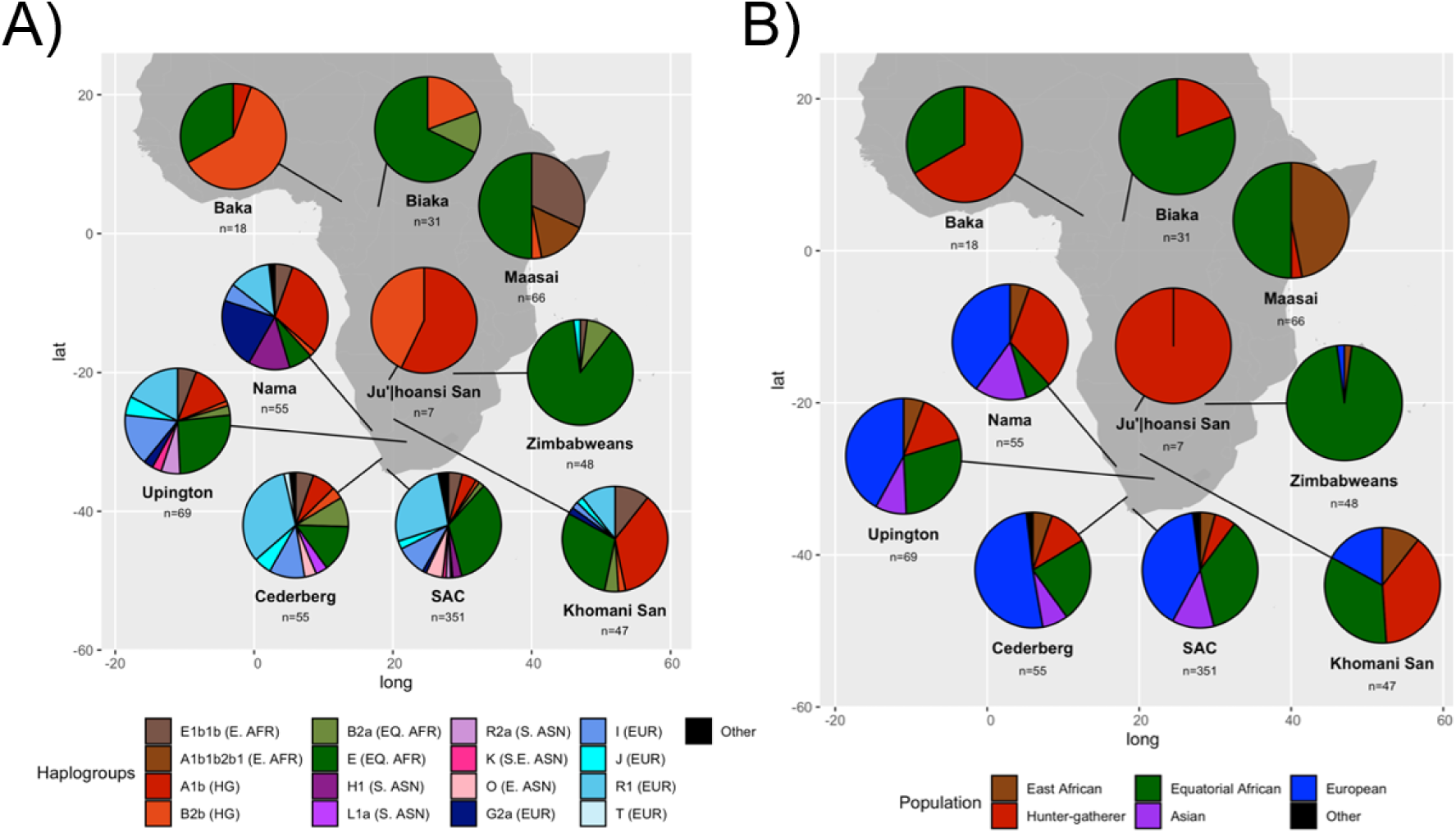
Distribution of Y chromosome haplogroups across southern Africa. A) The left panel shows the proportions of Y haplogroups present in samples from 10 different African populations, including five populations from South Africa, with comparative samples from Namibia, Zimbabwe, Kenya and Central African foragers. High-level Y haplogroups were used to assign individuals to a set of ancestries including: E.AFR=East African, HG=Hunter-gatherer, EQ.AFR=Equatorial African, EUR=European, ASN=Asian. “Other” includes 3 inconclusive assignments and five haplogroup assignments corresponding to only one individual (A0a1a [SAC], Q1a2 [SAC], C2e1b1 [SAC], S [Nama], E1b1b1b2a1 [SAC]). B) Broad ancestral origins for each of the Y haplogroups are displayed in order to summarize continental and sub-continental patterns for each population.

In South African populations, haplogroups endemic to the Khoe-San and Central African hunter-gatherers comprise a larger proportion of ancestry as one moves northward, from Cape Town to the Kalahari. The replacement of hunter-gatherer descendant Y haplogroups occurs with greater proximity to initial European colonial settlements in the Cape of Good Hope. Today, A1b and B2b haplogroups constitute only 6% of the Cape Town SAC lineages, the smallest proportion of such hunter-gatherer haplogroups across all southern African populations examined here (Fig. 4). Coloured populations from the Cederberg and Upington have 11% and 14.5% hunter-gatherer descendant haplogroups, respectively. This fraction increases appreciably in self-identified Khoe-San populations like the Nama (33%) and ≠Khomani San (38%). The Upington, ≠Khomani San and Nama hunter-gatherer haplogroups have the same proportional composition of 94% A1b and 6% B2b. The Cederberg hunter-gatherer haplogroup composition was slightly different at 66% A1b and 33% B2b.

We find evidence of major heterogenous male founder effects among different South African populations. There is a high proportion of European haplogroups, ranging from 40-51%, in all of the South African Khoe-San descent populations except for the ≠Khomani San. For example, among the 40% of the Nama Y-chromosomes that are of European descent; half of these are G2a2b, mostly G2a2b2a1a1b1a2b which is found infrequently across the continent. The Nama also have a sizeable proportion of Asian haplogroups (15%), almost all of which belong to the South Asian H1a1d2c1b1a haplogroup. Similarly, the Upington Coloured population has an enriched frequency of I2a2 lineages, which are rare enough to be absent among all 1000 Genomes Europeans, and 6% R2a South Asian lineages which are present in the Cape Town SAC at only 1.5% and otherwise absent in our sample. In contrast to the Nama, the ≠Khomani San have a much smaller proportion of European ancestry haplogroups (17%) and no Asian-derived haplogroups (Fig. 4). Finally, we observe unusual lineages like S, L1a, K2b, which derive from South Asia or Indonesian/Papua New Guinea, are non-uniform across the Coloured populations.

Interestingly, the E1b1b-M293 Eastern African haplogroup is found in all of the South African Coloured, Nama, and ≠Khomani San populations included here. This haplogroup is associated with the spread of pastoralism from Eastern Africa into Southern African hunter-gatherer populations (Bajić et al., 2018; Henn et al., 2008). The Nama and ≠Khomani San M293 sequenced Y-chromosomes all cluster in one monophyletic clade (Fig. S2), supporting a single pulse of male-biased migration into southern hunter-gatherer populations from Eastern African pastoralists 2,000 years ago (Vincente 2021). The Nama / ≠Khomani branch (along with a single Zimbabwean) is embedded in a cluster of Maasai M293 lineages. Other Kenyan and Tanzanian lineages form a sister clade to this cluster (Fig. S2). The frequency of M293 is twice as high in the ≠Khomani San (11%) relative to the Nama, Cederberg and Upington Coloured groups (5-6%). This observation is in conflict with previous autosomal data indicating a higher proportion of Eastern African ancestry in the Nama (7.4-8%) than ≠Khomani San (1.9%) (Vincente 2021). Vincente et al. (2021) propose that their results of decreasing Eastern African ancestry proportion in a North to South cline reflect the migration route of the pastoralists, with admixture dates becoming more recent the further south is sampled. While they showed that Eastern African ancestry was similarly male-biased in the Nama and ≠Khomani San (as well as among Hessequa and Khwe), our data suggest the dynamics of this migration deserve additional study.

### Timing of Male Population Growth and Decline

In order to explore the possible founder effects on the Y-chromosome, we estimate the male-specific demographic fluctuations of the three populations that contribute ancestry to contemporary Nama and ≠Khomani individuals using Bayesian Skyline Plots (BSP). After QC and removal of close paternal relatives, there were 89 individuals available for haplogroup assignment. The individuals were split into three groups corresponding to the population their Y-chromosome haplogroups affiliated with Khoe-San, European, or Equatorial African. Results of the BSP run for each population are shown in Fig. 5. We see a decline in the effective population size of Khoe-San male lineages around 120 generations ago (∼3,600ya), followed by a substantial recovery and growth. The effective population size of the Equatorial African lineages begins to expand shortly after 200 generations ago up to the present time. The European lineages in Nama and ≠Khomani individuals appear to be undergoing a strong bottleneck beginning around 20 generations ago.

**Fig. 5:**
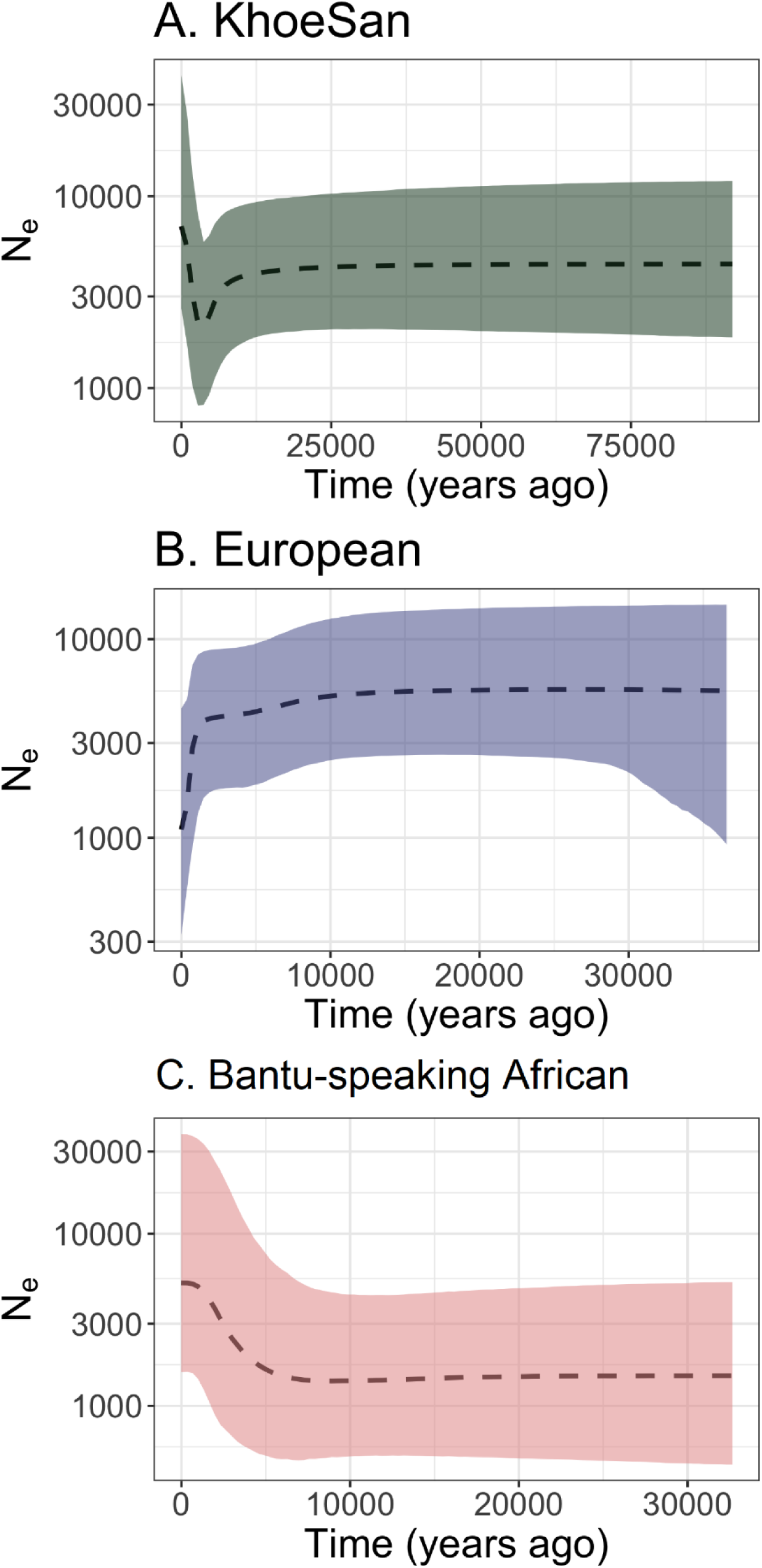
Bayesian Skyline Plots for Y chromosome haplogroups from the three major ancestry components. A) Khoe-San haplogroups (N = 39), B) Eurasian haplogroups (N=31), C) Bantu-speaking African (N=19). The dashed line is the median estimate of N_e_ and the color fill shows 95% HPD limits.

Based on the strong decline in N_e_ observed in the European lineages (Fig. 5B), we used coalescent simulations to determine if we could detect a demographic recovery following the colonial period with our data. Using *msprime* (Kelleher et al., 2016), we simulated 31 chromosomes under a demographic model similar to that observed in the empirical BSP (constant population size, followed by a steep bottleneck at 20 generations ago) with the addition of a demographic recovery at 10 generations ago. Our results (Fig. S3) show that Bayesian Skyline Plots would *not* be able to detect such a recent demographic recovery with data similar to the Y-chromosome data used here.

## Discussion

The effects of European colonialism are global and its legacy has impacted nearly every facet of society, including the genetic structure of indigenous populations. Using population genetic tools, we show the extent to which European male colonialists altered the population structure of indigenous southern African Khoe-San peoples. The mechanisms behind these colonial sex-biases are multi-faceted and include intermarriage as well as sexual violence (Penn 2014). This is clear from the genetic admixture seen in the Cape Town Coloured population, where only 6% of their Y-chromosomes has indigenous ancestry contrasted with 28% of their autosome, demonstrating a clear European male-bias. Like other indigenous populations around the world, Khoekhoe and San first contact with colonialists was shortly followed by dramatic population declines, from war/genocide and/or transmission of novel diseases, such as smallpox (Penn 2005). The settlement that would become the Cape Colony was a small supply station, with a focus on trading for livestock with the local Khoekhoe groups in the first three decades of its founding in 1652 (Guelke 1976). The settlement began to expand in 1679 to meet an increased need for provisions. During this period the European population of the Cape Colony expanded tenfold (from ∼200 to 2000 individuals). The increased need for labor was largely met by the VOC Indian Ocean slave trade, bringing individuals of both equatorial African (largely from Mozambique, Zanzibar, Angola, Madagascar) and Asian (largely from Indonesia and India) ancestry to the Cape (de Wit et al., 2010; Du Plessis, 1947).

In this context, we ask whether admixture among different ancestral groups occurred simultaneously or in a staggered fashion, and the extent to which differential sex-bias contributed to the SAC population in Cape Town. Our global ancestry results show strong patterns of male-biased admixture from Europeans and equatorial African populations, and correspondingly female-biased contributions from Khoe-San and Asian source populations. Our best-fit model infers three pulses of admixture of roughly equal proportions from all five ancestries, consistent with a model of continuous gene flow into the Cape Town SAC population. The formation of this population however is at odds with the historical record; our model estimates simultaneous contribution of all ancestries 22*ga* (approximately 1350-1550AD). While European explorers periodically interacted with Khoekhoe groups at the Cape throughout the 16^th^ century, e.g., kidnapping two men for an extended voyage to England (Kaufman 2017), a permanent settlement by the Dutch East India Company only occurred in 1652. Tract length distributions in our dataset include many short segments which may be poorly called, resulting in erroneous early time estimates.

### Expansion of the Colonial Frontier

As the Cape Colony expanded in the 18^th^ and 19^th^ centuries, new multi-descent ethnicities formed as the result of cultural interaction and intermarriage of Khoe-San, European, and enslaved Asian and equatorial African descent individuals in the Cape (Smith 1995). Their social standing in colonial society varied, but as the 18^th^ century progressed, discrimination towards so-called “Bastaard” communities increased, resulting in individuals moving into the northern and eastern frontiers of the Cape Colony. As these individuals fled from the expanding colony, they encountered other indigenous KhoeSan groups and in many cases became integrated with them, with recorded examples in the Nama (Penn, 2005), or formed entirely new communities (Edgar and Saunders, 1982; Nurse and Jenkins, 1975; Ross, 1975). As the frontier continued to expand and the Dutch established missions and permanent settlements further north, Khoe-San groups coalesced near the Orange River as their social structures were disrupted by disease and pressure to become a part of the colonial culture (Smith 1995).

By examining the distribution of Y chromosome haplogroups, we can infer the ancestry of individuals moving into the periphery of the Cape Colony. The major haplogroup ancestry proportions in the Cape Town SAC are remarkably similar to the proportions in two other SAC populations in the Cederberg Mountains and the Orange River (Fig. 4). All groups retain ∼10-20% indigenous E1b1b, A1b, and B2b haplogroups indicative of the Khoe-San populations. European-derived haplogroups constitute 40-50% of all men. These Y-chr proportions stand in stark contrast to SAC mitochondrial ancestry, which Quintana-Murci *et al*. (2010) found to be only 4.6% European. The Asian Ychr haplogroups have the highest diversity in Cape Town, with subsequent drift further north, e.g. the loss of haplogroup O in the Northern Cape. These patterns suggest a model of relatively large-scale migration from the Cape into the colonial frontier, and the establishment of acculturated mixed communities. It remains to be tested whether these communities absorbed additional input from local Khoe-San groups and whether gene flow from the European men was a continuous process in the divided society. In contrast, Y haplogroups between the Nama and ≠Khomani San are quite distinct to the SAC, while still exhibiting strong sex-biased impacts.

### Richtersveld Nama

Nama, a group of Khoekhoe pastoralists, have lived in the Richtersveld region for hundreds of years prior to the expansion of the Cape Colony (Hoernle 1913; Carstens 1985). Here, we show that the Richtersveld Nama exhibit ∼29% genome-wide European ancestry, and negligible Asian ancestry compared to the SAC from Cape Town. The highest likelihood model for tempo of admixture suggests that there were three gene flow events in the Nama. The first represents a gene flow event between Khoe-San and equatorial African ancestry ∼28 generations ago. This is likely due to interactions between the nothern Namibian Damara who are of Western African ancestry (Barbieri et al., 2014; Montinaro et al., 2017). The second and third pulses of gene flow occur in quick succession, with the Nama first receiving ∼30% ancestry from Europeans approximately 7-8 generations ago (210-240 years ago assuming a generation time of 30 years) followed by gene flow from Khoe-San 6-7 generations ago. This is consistent with the patterns seen in the ≠Khomani San (Uren et al. 2016) and potentially indicates the dynamics of Khoe-San retreating from the colonial frontier. This process may have resulted in reorganization of fine-scale population structure and the merging of multiple groups.

Y-chromosome haplogroups provide greater resolution for the sources of colonial migration into the Nama. Just under half of the Richtersveld Nama have European-derived haplogroups, and 15% derive ancestry from South Asia. Several of these haplogroups display evidence of founder effects, falling into single, highly derived clades such as G2a2b2a1a1b1a2b and H1a1d2c1b1a. Such strong founder effects supports a model of infrequent outside contribution to the Nama by only a few men, but disproportionate reproductive success. Cultural connections -- such as speaking Afrikaans -- could have facilitated access to trade or other resources; alternatively, these men may have been viewed as of higher status and disproportionately desirable as partners.

The demographic reconstruction based on Y chromosome capture data in the Nama and ≠Khomani provide further insights into the male population size changes during the colonial period. Unexpectedly, we find that the European Ychr lineages in our dataset underwent a recent severe bottleneck, but a recent study of the South African Afrikaner population found no evidence of a genome-wide bottleneck in this European-descendant group (Hollfelder et al., 2020). Simulations (Fig. S3) suggest that the Y chromosome Bayesian Skyline pattern is consistent with a founder effect among European lineages around the time of initial colonization of South Africa. A very recent population recovery would not alter the pattern seen in the BSP.

We examined the spatial and temporal spread of sex-biased migration from the Cape northward into Namaqualand and the southern Kalahari using genetic data from more than 1,400 individuals. In all regions, admixture patterns were sex-biased, with evidence of a greater male (compared to female) contribution of European ancestry, and greater female contribution of Khoe-San ancestry. These genetic data show the direct population impact of European colonialism on population structure in southern Africa. The extent to which local indigenous Khoe-San populations were replaced or integrated into mixed communities, and the mechanisms of sex-bias remain critical open questions for future study.

## Materials and Methods

### Samples and Data

#### SAC Data

Study participants were recruited from two suburbs in the Cape Town Metropol in the Western Cape, for the purpose of understanding tuberculosis susceptibility (1995-2005). DNA was extracted from blood samples of 800 individuals, who self-identify as SAC, as reported previously (Schurz et al., 2018). DNA extracts from the SAC individuals were genotyped for ∼1.6 million SNPs on the Illumina Multi-ethnic genotyping array (MEGA) (Illumina, Miami, USA).

#### Nama and ≠Khomani San Data

Saliva samples were collected from 110 Nama individuals from the Richtersveld region of South Africa in 2014 and 2015. Saliva samples were collected from 170 ≠Khomani San individuals southern Kalahari Desert of South Africa in 2006, 2010, 2011, 2013, and 2015. Samples were collected with Oragene OGR-500 kits (DNA Genotek) and extracted using prep-IT L2P reagents (DNA Genotek) according to the manufacturer’s protocol. All samples were genotyped for ∼1.6 million SNPs on the MEGA array (Illumina, Miami, USA).

#### Ethics Approval and Return of Results

DNA samples from the Nama, ≠Khomani San, Cederberg, NCTB and SAC populations were collected with written and informed consent. Institutional review board (IRB) approval was obtained from Stanford University (protocol 13829), Stony Brook University, University of California at Davis, and the Health Research Ethics Committee of Stellenbosch University (N11/07/210, N11/07/210A, N95/072, S17/01/013 and N06/07/132), South Africa. A contract for this project was approved by the Working Group of Indigenous Minorities in Southern Africa. Local community members, local community leaders, traditional leaders, nonprofit organizations, and a legal counselor were all consulted regarding the aims of the research before collection of DNA.

#### Genotype Array QC

Raw genotype data was QC’d using Illumina’s GenomeStudio to call common variants (MAF>0.05), followed by zCall to call rare variants (Goldstein et al., 2012). All datasets were aligned to the 1000 Genomes Phase 3 reference panel and all ambiguous variants were removed. Plink v 1.07 (Purcell et al., 2007) was used to check for sex concordance prior to merging all datasets. The merged data was filtered for Hardy Weinberg Equilibrium (<0.05), minor allele frequency (<0.03) and individual and SNP missingness (>10%) using plink (v1.07). Finally, all variants on the X chromosome that were heterozygous in males were removed and the merged dataset was checked for ambiguous variants using plink (v1.07) and snpflip (v0.0.6, https://github.com/biocore-ntnu/snpflip) respectively. PAR regions of the X and Y chromosomes were removed for our analysis.

### Y-chromosome analysis

#### Y-chromosome Illumina capture

272 male samples were selected from the Nama and ≠Khomani San, as well as several comparative populations (Zimbabwe, Maasai, Baka, Biaka, Ju|’hoansi) underwent target capture for 8.9Mb of the Y-chromosome informative for detailed haplogroup assignments (Cruz-Dávalos et al., 2018; Poznik et al., 2013). Libraries were prepared (Bentley et al., 2008) and then sequenced the libraries on Illumina HiSeq 2000 machines at the Stanford Center for Genomics and Personalized Medicine. The samples had a mean of 10x coverage (range 1-70x) of the targeted regions of the Y-chromosome (Supp Table 1).

Raw FASTQ files for the forward and reverse strands were merged using leeHom (Renaud et al., 2014). Adapters were trimmed from the merged files using fast-qc (Andrews, S, 2010) and mapped to the GRCh37 human reference using BWA (Li and Durbin, 2009). Chimeric and low-quality read pairs were then removed from merged and paired-end alignment files. GATK v4 was then used for duplicate removal, local realignment for indels, and base quality recalibration. Finally, the merged and paired-end BAM files were merged, deduplicated, and filtered base on mapping quality (MQ > 40).

Filtered BAM files were then aligned and indexed using the SAMtools (v 1.3.2) -sort and index flags respectively (Li et al., 2009). Prior to variant calling duplicated reads were removed using picard tools (v 2.23.4) MarkDuplicates and REMOVE_DUPLICATES flags, followed by re-indexing of the file using SAMtools. In preparation for GATK, picard tools AddOrReplaceReadGroups option was used to replace read groups and update the platform information to Illumina, after which the file was again indexed using SAMtools.

GATK v4.1.6.0 (McKenna et al., 2010) was used to call variants in the individual files using the human_g1k_v37.fasta and the 1000 Genomes Phase 3 data (The 1000 Genomes Project Consortium, 2015) as references. First bases were calibrated in each individual sequencing file using the GATK BaseRecalibrator and ApplyBQSR commands. Next individual haplotypes were called by implementing the GATK HaplotypeCaller option and as we are analysing Y chromosome data the ploidy was set to 1 (-ploidy 1) for haploid data analysis. Individual GVCF files with called haplotypes were then merged into one GVCF file via the GATK CombineGVCFs command. Variants for all samples were then recalled jointly following GATK best practices (Van der Auwera et al., 2013), using the GATK GenotypeGVCFs (again with the -ploidy 1 flag) and SelectVariants (-select-type SNP flag) commands.

The selected variants were then inspected for quality and marked for removal if they had a quality by depth score less than 2, mapping quality score less than 30, Fisher strand score greater than 60, mapping quality rank sum test value less than -12.5, or read position rank sum score less than -8.0 via the GATK VariantFiltration option (--filter-expression "QD<2.0 || MQ<30.0 || FS>60.0 || MQRankSum<-12.5 || ReadPosRankSum<-8.0"). Variants that failed any of the criteria were then removed and filtered genotype sites were set to missing using the GATK SelectVariants option and –set-filtered-gt-to-nocall flag.

The filtered file was then condensed and indexed using bgzip and tabix (Li, 2011) respectively and then a bed mask was applied to remove off target regions using bcftools v1.12 (Danecek et al., 2021). Once these sites were removed bcftools was implemented to obtain the read depths of the called variants (query -f ’%CHROM %POS %DP\n’), which were then plotted in R in order to determine valid minimum and maximum cut-off values.

Finally, GATK VariantFiltration was again implemented to mark variants that were above or below 3 MADs of the median depth value (--filter-expression "DP<1515 || DP>9557"), these were then removed using the GATK SelectVariants option to obtain the final VCF file for the Imputation step. Four samples were removed due to low mapping quality prior to imputation.

#### Y chromosome Imputation

To impute missing genotypes in the called and filtered file we implemented Imputor (Jobin et al., 2018) an imputation software specifically designed to impute haploid non-recombining loci, such as the human Y-chromosome. Prior to imputation, individuals with more than 25% missing genotypes were removed using vcftools v1.17 (Danecek et al., 2011). Three passes of Imputor were then run on the dataset using the hops algorithm, which imputes missing sites using the nearest neighbors determined by the total number of “hops” from node to node along the tree (Jobin et al., 2018). The resulting imputed dataset was again filtered for individual missingness of >10% and genotype missingness of >5% using plink v1.9 (Chang et al., 2015, p. 9) to obtain the final file to be used for downstream analysis. Seven samples were removed due to missing genotypes and an additional 8 samples were removed for having a close male relative in the dataset, resulting in a final Y-NGS sample size of 89 individuals in the Bayesian Skyline dataset.

#### Y chromosome haplogroup assignment

Y chromosome haplogroups were called using two different software packages. SNAPPY (version 0.2.1) was used to assign haplogroups for the genotype array data, as it is optimized for use with the Illumina MEGA array that the SAC (Cape Town) are genotyped on as well as the Illumina H3Afriaca array that the Cederberg and Upington samples are genotyped on (Severson et al., 2018). Haplogroups were assigned for Y chromosome data using the yhaplo software (Poznik, 2016).

### Admixture inference

#### Global Ancestry

For the admixture analysis the software ADMIXTURE 1.3 was used (Alexander et al., 2009). The reference populations used to infer ancestry were Europeans (Utah Residents with Northern and Western European Ancestry) and South Asian (Gujarati Indians in Houston, Texas and Pathan of Punjab) obtained from the 1000 Genomes Phase 3 data (Sudmant et al., 2015) and East Asian (Han Chinese in Beijing, China and Japanese, Tokyo, Japan), African (Luhya in Webuye, Kenya, Bantu-speaking African, Yoruba from Nigeria) and KhoeSan (Nama and ≠Khomani) (Martin et al., 2017; Uren et al., 2016). Due to the level of relatedness in the SAC and the limited number of individuals per reference population the SAC individuals were split into 20 running groups with an average of 42 unrelated SAC individuals per group, matching the number of individuals per reference population. Relatedness for the SAC was determined from the genotyping data using the software KING (version 2.1.4) (Manichaikul et al., 2010). For each running group admixture was inferred five times at random seed values for both the autosome and X chromosome separately. For the X chromosome the ADMIXTURE software was run in haploid mode for males (--haploid=”male:23”) in order to ensure accurate admixture inference for haploid genotypes (Shringarpure et al., 2016). The values for the five runs were then averaged for each individual before the results were analysed. Admixture was run at K=5 as we were interested in the 5 main SAC ancestral components. The reference populations used to infer ancestry were European (CEU) and South Asian (Gujarati Indians in Houston, Texas and Pathan of Punjab) extracted from the 1000 Genomes Phase 3 data (Sudmant et al., 2015), East Asian (Han Chinese in Beijing, China), African (Luhya in Webuye, Kenya, Bantu-speaking African, Yoruba from Nigeria) and Khoe-San (Nama and ≠Khomani) (Martin et al., 2017; Uren et al., 2016). Pong (version 1.4.7) was used to visualise the admixture results across all runs (Behr et al., 2016).

Admixture inference for the Nama and ≠Khomani data was done in the same way, except that K was set to three as these are less admixed populations. Reference populations used for both Nama and ≠Khomani were from the African (Luhya in Webuye, Kenya, Bantu-speaking African, Yoruba from Nigeria) and European (French, Orcadian) populations (Martin et al., 2017; Uren et al., 2016). For the Khoe-San ancestry in the Nama, the ≠Khomani were used as a reference and for the ≠Khomani we used the Nama population as a reference to infer the Khoe-San ancestral component. Due to the relatedness in the Nama and ≠Khomani samples the data was split into running groups of unrelated individuals for inference of ancestry. Nama and ≠Khomani data were split into 7 and 8 running groups respectively and relatedness was again determined using KING software (version 2.1.4) (Manichaikul et al., 2010).

#### Local Ancestry Assignment

##### Phasing

Phasing of the Nama and SAC data was done using ShapeIT2 (Delaneau et al., 2013) and the same reference populations used for the global admixture inference. References and population of interest (Nama or SAC data) were merged into one file for local ancestry inference. For the autosome the data was split into individual chromosomes prior to phasing, using default ShapeIT2 settings and implementing the --duohmm option in order to take relatedness into account when inferring local ancestry. Phasing for the X chromosome was done slightly differently. First males and females were phased together in order to fill in missing male data based on the female data. Following this the phased files were split into males and female files and then the females were rephrased using the males as a reference. As males have no X chromosome recombination and thus the entire chromosome is one haplogroup they serve as a good reference to increase accuracy of local ancestry inference in females.

##### RFMix

RFMix v1.5.4 (Maples et al., 2013) was used to infer local ancestry using the phased per chromosome files form ShapeIT2 containing both population of interest and references. Local ancestry was inferred for each autosomal chromosome separately using default RFMix settings except that –G was set to 14, -e was set to 3, -n was set to 5 and RFMix was set to reanalyse references in order to improve accuracy of local ancestry inference. For the X chromosome the input files were split based on sex and inference was done separately, using the same settings as for the autosome except that the chromosome was set to 23 to inform RFMix that this is a sex specific chromosome. The genetic map used was the same one that was used in the Phasing step except it has been edited to work with RFMix.

### Sex bias analysis

#### Global Admixture

Sex bias was analysed by comparing the distribution of each ancestral component between the autosomes (1-22) and X chromosome. The data was assessed for normality (Fig. S1) upon which a paired Wilcoxon signed rank test (for skewed data) was implemented, using the R programming environment (version 3.2.4 (R Development Core Team, 2013)), to determine significant differences in ancestral distribution. If the mean ancestral component is significantly larger on the autosome compared to the X chromosome, then it is an indication of male bias, while a larger X chromosomal mean is indicative of a female bias. To minimise the multiple testing burden (Bonferroni correction, alpha threshold of 0.05) we limited the number of ancestral components to 5 for the SAC and 3 for the Nama population. While previous studies have suggested higher levels of admixture for these populations, small ancestral components combined with increased multiple testing burden is unlikely to produce significant results for the sex bias analysis (Uren et al., 2016).

### TRACTs analysis

To estimate the time intervals of admixture for the Nama and SAC individuals we modelled historical time-dependent gene-flow from the various source populations using TRACTs (Gravel, 2012). For both the Nama and SAC we evaluated several two, three and four-pulse models, ordering the populations as KhoeSan, Bantu, European for Nama and KhoeSan, Bantu, European, Asian for the SAC. Source populations for ancestral haplotypes includes: KhoeSan (represented here by Nama / ≠Khomani), equatorial African (represented here by the Kenyan Luhya and Nigerian Yoruba), European (represented here by the CEU, Central European) and Asian (represented here by Gujarati Indians, Han Chinese and Japanese). Analysis of the 5-way admixed Sac population required adaptation of the TRACTs scripts to include a fourth ancestry and tested 15 models (Table S1). The Asian component of the SAC is a combination of South and East Asian ancestry that was grouped together into Asian ancestry as the small contribution from East Asia would introduce errors into the TRACTs analysis. For the Nama individuals we tested the same 9 models (Table S1) previously implemented for the ≠Khomani San population (Uren et al., 2016). Each model (for Nama and SAC separately) was tested three times and then the top three models were further evaluated by fitting it with 100 randomised starting parameters and the best fit model will be the one with the lowest log-likelihood value. In Table S1 and S2 each letter (x or p) corresponds to the population order given above. The ‘p’ presents a pulse and the ‘x’ no pulse from the corresponding ordered ancestry and the underscore ‘_’ indicates a migration event.

### BEAST

In order to determine male-specific population growth within the KhoeSan, we applied a Coalescent Bayesian Skyline model implemented in BEAST v 2.6.2 (Suchard et al., 2018) to the NGS Y-chromosome dataset. We assume a general time reversible (*GTR*) substitution model with 4 gamma categories to allow for transition and transversion rates to vary. The analysis was performed using an uncorrelated exponential related (*UER*) clock with a constant rate of 0.82e-9 (Poznik et al., 2013). Y-chromosomes were assigned to 3 different ancestries based on their haplogroup affiliation: Eurasian, KhoeSan and Bantu-speaking African – and then each subset analysed separately. Imputed fasta files were loaded into BEUti version 2 to set variables and implement the ECBS analysis for the BEAST run. Four independent runs were performed for each group with a chain of 100 million iterations. Parameters were logged every 10,000 steps. The resulting log and tree files were combined using BEAST’s logCombiner, and a 10% burn-in was removed from the combined files. BEAST output files were analysed using Tracer (v 1.7.1) (Rambaut et al., 2018) and R to plot the skyline plot and determine growth over time for the specific sub-populations.

#### European demographic simulations

We used msprime v0.7.6 (Kelleher et al., 2016) to simulate Y-chromosome data under a demographic model consistent with what is known about the colonial history of European males in South Africa. We simulated genetic variation along a chromosome of the same length and average mutation rate as the human Y-chromosome. Our empirical model included a bottleneck consistent with the timing of the Out-of-Africa event ∼2400 generations ago (Gutenkunst et al., 2009), a long period of bottleneck at 20 generations ago, and a recovery starting 10 generations ago. We did this for 31 independent chromosomes to match our empirical dataset. To best emulate the empirical dataset, we randomly subsampled the simulated chromosomes to the number of variants in the empirical dataset. Because variant information in msprime is binary, we also matched the transition/transversion ratio found in the empirical dataset. The simulated data was then run through BEAST in the same manner as the empirical data.

## Supporting information captions

**Table S1:** TRACTs result for the different models in the SAC population with 4 ancestral components (population order: San, Bantu, European, Asian)

| Model | Likelihood |
| --- | --- |
| ppxx_xxpp | -3166.95 |
| ppxp_xpxx | -2956.33 |
| ppxx_pppp | -3085.67 |
| pppp_xppp | -1783.76 |
| pppp_xpxx | -2318.87 |
| pppp_pxxx | -2806.91 |
| ppxx_xppp | -1783.76 |
| pppx_xxpp | -3176.72 |
| pppp_pppp | -738.03 |
| ppxx_xppx_xxpp | -2167.17 |
| ppxx_xpxx_xxpp | -3305.32 |
| ppxx_xxpp_xpxx | -3306.06 |
| ppxp_xpxx_pxxx | -2486.33 |
| ppxx_xxpp_pxxx | -2767.85 |
| ppxx_xppp_xpxx | -2588.95 |
| ppxx_xxpp_xxpp | -3014.59 |
| ppxx_xppp_xxpp | -2934.46 |
| pppp_xxpp_xpxx | -2805.77 |
| pppp_xxpp_ppxx | -1906.23 |
| pppp_xxpp_xxpp | -3255.38 |
| pppp_xppp_ppxx | -784.58 |
| pppp_pppp_xpxx | -2830.47 |
| pppp_pppp_ppxx | -576.23* |
| **pppp_pppp_pppp** | **-540.73\*** |
| pppp_xppp_ppxx_xpxx | -2377.07 |
| pppp_pppp_ppxx_xpxx | -719.47 |
| pppp_xppp_xxpp_ppxx | -796.01 |
| pppp_xpxx_xppp_pppp | -735.07 |
| pppp_xppp_pppp_xpxx | -753.73 |
| pppp_pppp_pppp_pppp | -833.56 |
| pppp_pxxx_xpxx_xpxx | -1203.15 |
| pppp_pxxx_xpxx_xxpp | -621.55 |
| pppp_pppp_pppp_xppp | -558.53* |

\* Top 3 most likely TRACTs models

**Table S2:** TRACTs result for the different models in the Nama population with 3 ancestral components (population order: San, Bantu, European)

| Model | Likelihood |
| --- | --- |
| ppx_xxp | -640.9 |
| ppp_xxp | -533.5 |
| ppp_pxp | -486.90* |
| ppp_ppp | -759.71 |
| ppx_xxp_pxx | -648.602 |
| ppx_xxp_xpx | -737.20 |
| ppx_xxp_pxp | -598.92 |
| **ppx_xxp_ppx** | **-432.51\*** |
| ppx_xxp_xxp | -615.07 |
| ppp_xxp_pxx | -454.20* |

\* Top 3 most likely TRACTs models

**Table S3:** Y chromosome haplogroup assignments by population.

**Figure S1:**
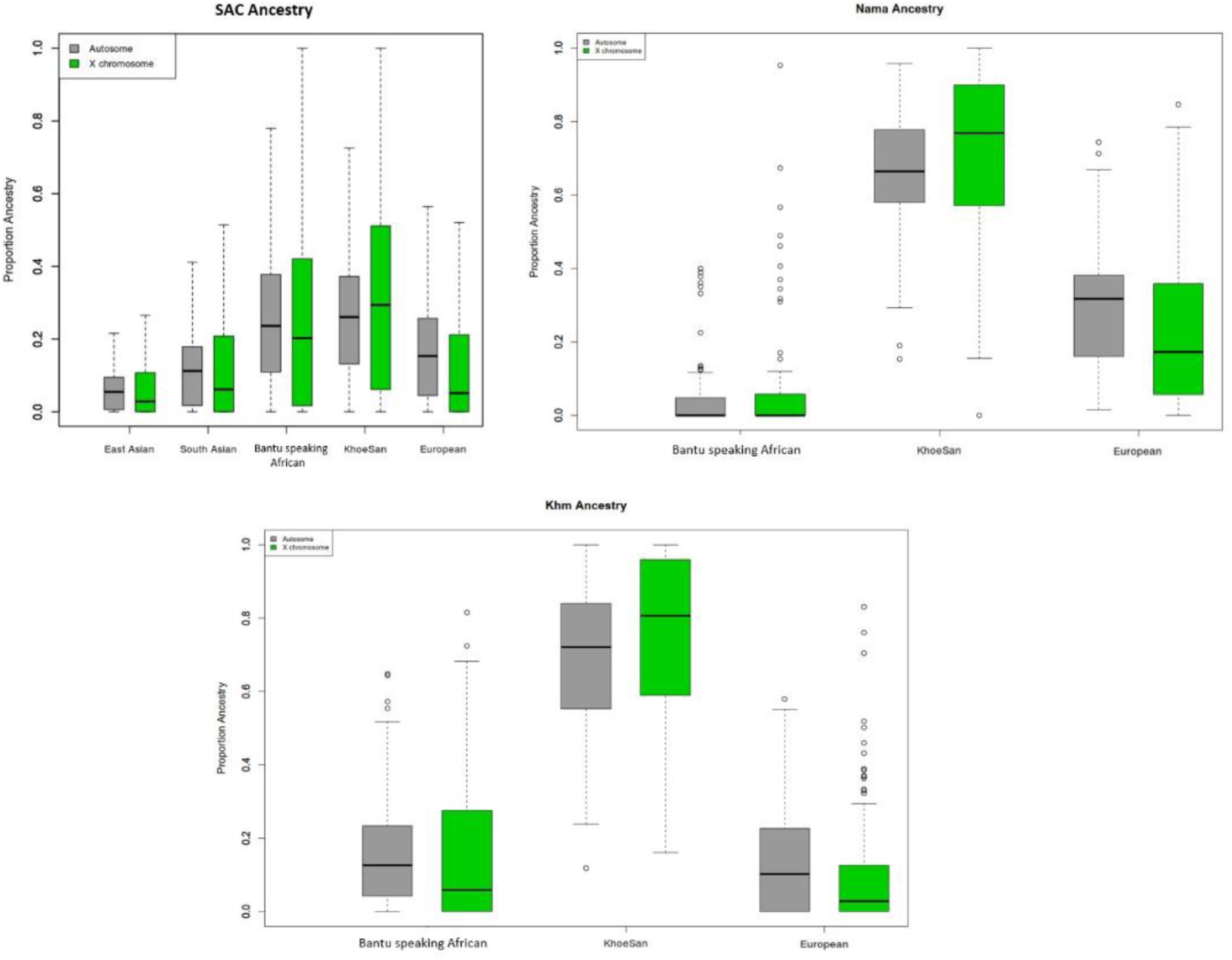
Normality assessment for admixture components.

**Figure S2:**
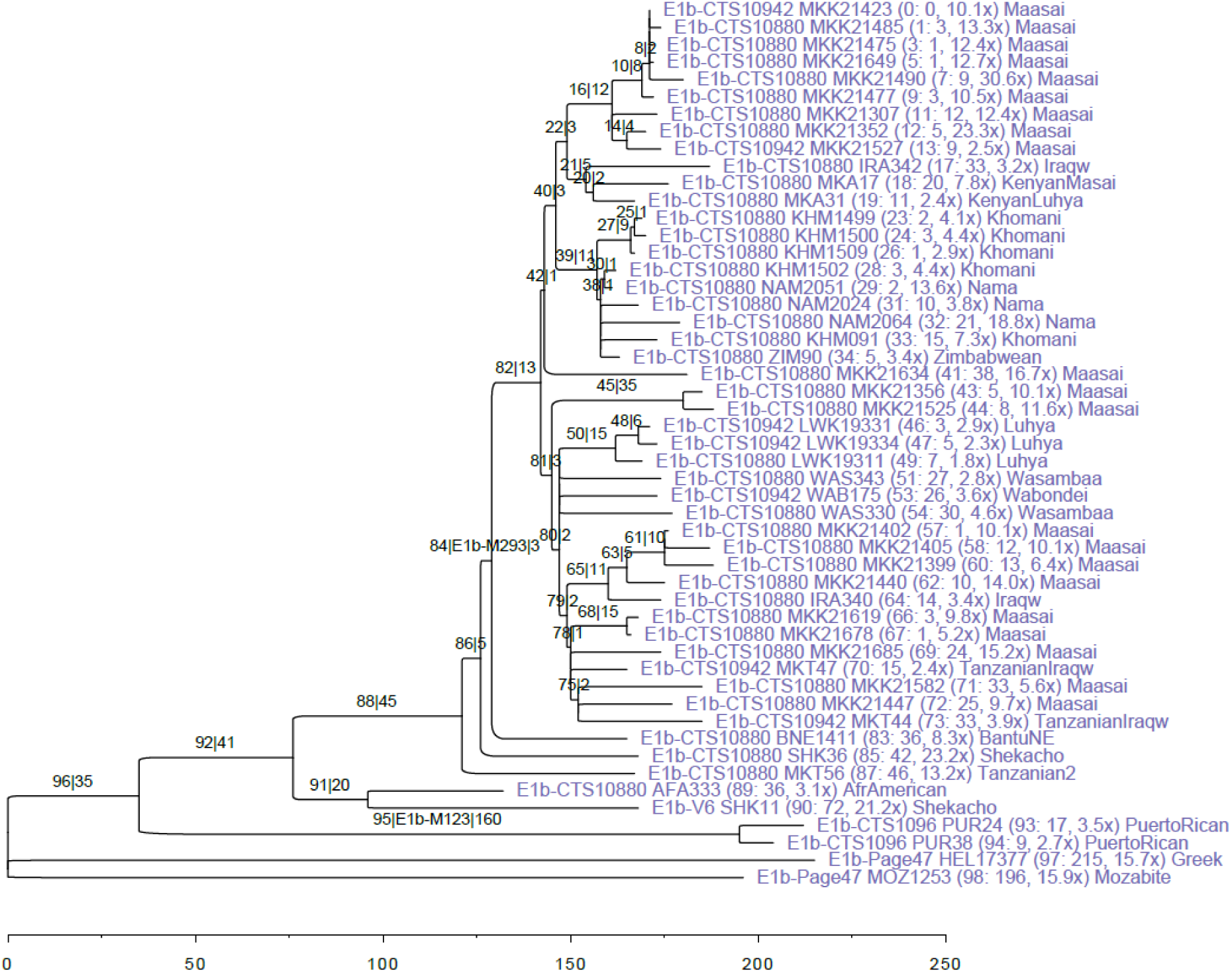
Maximum likelihood tree for Y-chromosome haplogroup E1b1b-M293 constructed with RAxML. Branch lengths are drawn in proportion to the number of SNPs that map compatibly to each branch. Internal branches are labeled with an index, a canonical SNP (major branches only), and the branch length, separated by pipes (|). Terminal branches are labeled with the individual’s haplogroup, most derived ISOGG SNP, and sample ID, then, in parentheses, the branch index (followed by a colon), branch length (number of singletons), sequencing coverage, and finally the population.

**Figure S3:**
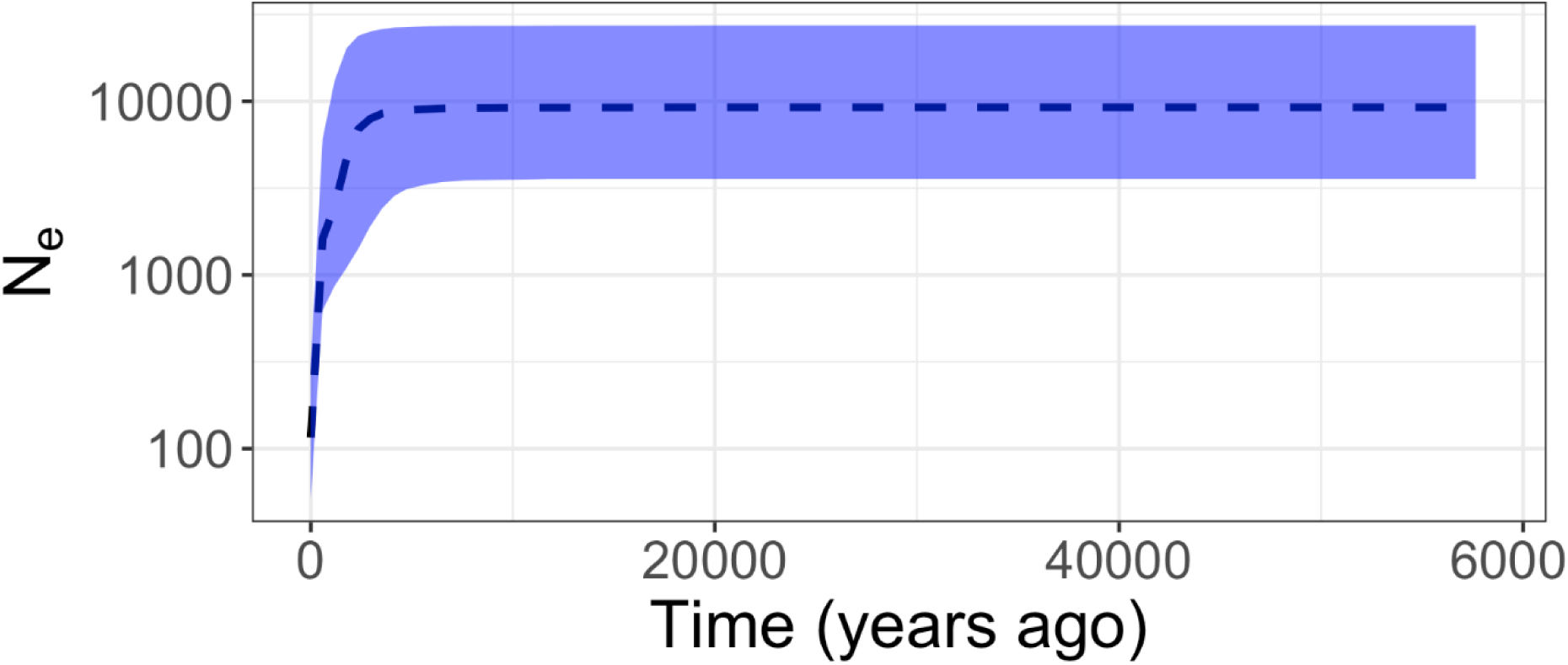
Bayesian Skyline plot based on data simulated under a demographic model of constant population size of 10000 individuals followed by a bottleneck of 100 individuals 20 generations ago and a full recovery 10 generations ago, consistent with historic records of the expansion of European colonists in South Africa.

